# CODEX, a neural network approach to explore signaling dynamics landscapes

**DOI:** 10.1101/2020.08.05.237842

**Authors:** Marc-Antoine Jacques, Maciej Dobrzyński, Paolo Armando Gagliardi, Raphael Sznitman, Olivier Pertz

## Abstract

Fluorescent biosensors routinely yield thousands of single-cell, heterogeneous, multi-dimensional signaling trajectories that are difficult to mine for relevant information. We present CODEX, an approach based on artificial neural networks to guide exploration of time-series datasets and to identify motifs in dynamic signaling states.

---

Dynamic rather than static signaling states control fate decisions^1^. Biosensor imaging has documented heterogeneous and asynchronous p53^2^, NF-kB^3^, ERK^4,5^, Akt^6^, SMAD^7^ dynamic signaling states across individual cells of a population. This heterogeneity arises due to both biological noise extrinsic to individual cells and intrinsic variability within signaling networks^8^. Current biosensor measurements yield up to thousands of heterogeneous, potentially multivariate, signaling trajectories^6^. The characterization of these trajectories typically relies on visual inspection, followed by statistical analysis of extracted features designed to be human interpretable^6,9^. The volume and high dimensionality of these datasets, as well as the capacity of features to faithfully describe complex characteristics of the trajectories, make the mining challenging. To meet these challenges, we present CODEX (COnvolutional neural networks for Dynamics EXploration), a data-driven approach for the exploration of dynamic signaling trajectories using convolutional neural networks (CNNs). The approach benefits from the ability of CNN classifiers to identify a set of data-driven features that best summarize the data.

In a typical CODEX analysis, a CNN classifier is trained to recognize specific cellular states (the input classes) based on their corresponding signaling trajectories (Fig.1a). To circumvent the difficulties related to CNN training, we use a plain, yet powerful CNN architecture (Supplementary Note 1, Table S1). We first benchmarked CODEX using synthetic trajectories that mimic pulsatile signaling states typically observed for different pathways^5,10^ (see Methods and Supplementary Fig.S1, Supplementary Note 2). This revealed that the features learnt by the CNN recognize discriminative features of the dynamics within the input. A t-Distributed Stochastic Neighbor Embedding (t-SNE) projection of the CNN features revealed discernible clusters that corresponded to specific dynamic patterns (Supplementary Fig.S1a,b). We then showed that this feature space could be sampled to identify different prototype trajectories representative of all dynamic states (Supplementary Fig.S1b). Further, we used Class-Activation Maps (CAMs)^11^, a technique that reveals parts of the input that draw the CNN’s attention towards a given class, to identify intervals within trajectories that were specific for a class (Supplementary Fig.S1c, Supplementary Video 1). Then, we clustered these intervals using dynamic time warping (DTW) distance to reveal groups of class-specific dynamic motifs (see Methods and Supplementary Fig.S1d).

**Figure 1:**
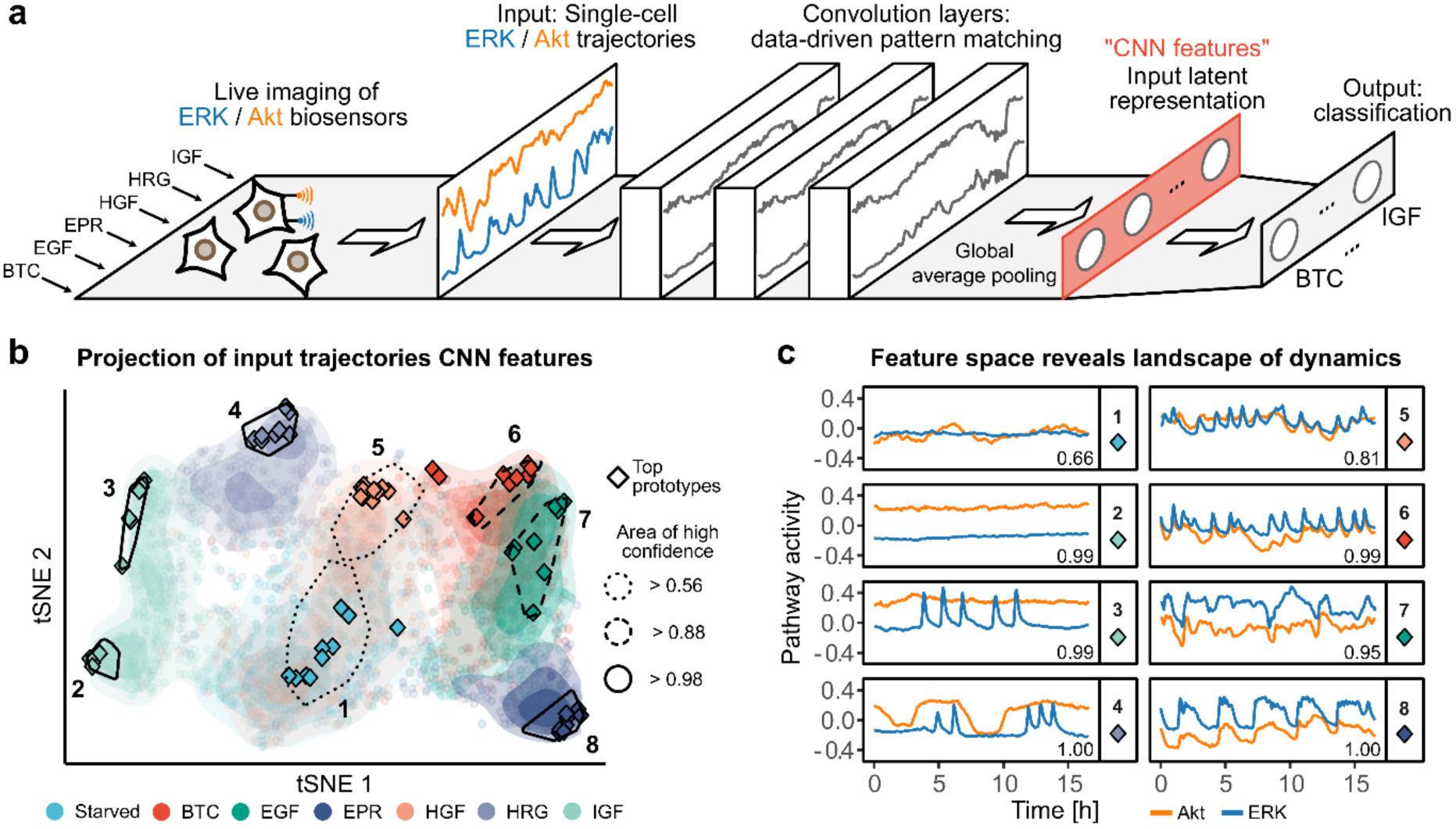
CNN latent features reveal the landscape of single-cell signaling dynamics. **a**, Schematic of the CNN classifier architecture used in CODEX. In this example, MCF10A cells are exposed to 6 different GFs and single-cell ERK/Akt activity is reported using biosensors. The GF treatments form classes that the classifier is trained to identify based on the bivariate ERK/Akt single-cell input trajectories. The input trajectories are passed through a cascade of convolution layers, followed by a global average pooling layer that compresses the convolution responses into a one-dimensional latent features vector (red rectangle). This latent representation of the input is then referred to as CNN features. Finally, the CNN features are passed to a fully connected layer to perform the classification. **b**, t-SNE projection of the CNN features of all ERK/Akt trajectories in the validation set. Each point represents a bivariate ERK/Akt trajectory from a single cell. Hulls indicate areas associated with a strong classification confidence for each GF. Shading shows the point densities. Diamonds indicate the positions of the 10 top prototypes (see Methods) for each GF. The solidity of the hull contour line indicates the minimal classification confidence for all prototypes in the hull. **c**, Representative ERK/Akt prototype trajectories from each hull indicated in (**b**). Each pathway activity is preprocessed by removing its respective average in the training set. Numbers in the bottom-right corners indicate the CNN predicted probability for the trajectories to belong to their actual class.

After this proof of concept, we applied CODEX to study Ras-ERK/PI3K-Akt single-cell signaling dynamics that was measured using a multiplexed genetically-encoded biosensor system, and quantified using a computer vision pipeline (Supplementary Fig.S2a,b). We stimulated MCF10A breast epithelial cells with different growth factors (GFs) known to induce distinct dynamic ERK/Akt signaling states^6^. We used 100 ng/ml of: epidermal GF (EGF), betacellulin (BTC), epiregulin (EPR), hepatocyte GF (HGF), heregulinβ-1 (HRG) and insulin-like GF 1 (IGF); an additional class consisted of starved, untreated cells. For each GF, we acquired at least 1200 of single-cell, bivariate ERK/Akt trajectories across two replicates. In the 1st phase, that lasted about 8 hours, population-synchronous signaling states were observed due to acute GF stimulation (Supplementary Fig.S2c–e). This is followed by the 2nd phase which is characterized by heterogeneous signaling states across the cell population. These states are relevant for proliferation fate decisions^5,6^. We truncated the trajectories to keep the 2nd phase only. To explore the specific ERK/Akt states induced by each GFs, we trained a CNN classifier that takes bivariate ERK/Akt trajectories as input to predict starvation or treatment with a specific GF. After training, this classifier separated the different classes with about 65% accuracy (Supplementary Table S2, S3, suggesting that input trajectories carry distinctive features that depend on GF identity.

To map the landscape of signaling dynamics, we used t-SNE projection of the CNN features learnt for each trajectory (Fig.1b). We observed that the GFs populated well-defined but also slightly overlapping areas of the projection. Trajectories of starved cells localized to a central, largely spread cluster (area 1). West of area 1, IGF (IGFR ligand) led to a polarized cluster that we arbitrarily designated as areas 2 and 3, while HRG (ErbB3/4 ligand) formed a distinct cloud (area 4). HGF formed area 5 that overlapped greatly with starved cells. East of area 1, a group was formed by EGF, EPR and BTC which are all ErbB1 ligands. This suggests that each GF induces a specific continuum of heterogeneous signaling states, whose characteristics correlate with their cognate GF receptor. Finally, we noted that GFs which overlapped in the projection, corresponded to cases where the CNN confusion was high (Supplementary Table S2).

To interactively explore this space, we devised a software to visualize the trajectories in different areas and report their associated ERK/Akt states (Fig.1b, Supplementary Video 1). We first evaluated trajectories that exhibited highest classification accuracies for each class, that we refer to as “top prototypes” (see Methods, Fig.1c, Supplementary Fig.S3). Through visual examination, we empirically describe the main trends of the prototypes using qualitative features. Starved cells displayed flat baseline ERK/Akt levels (Fig.1c, area 1). The polarized cluster induced by IGF revealed two distinct signaling states: both displayed sustained Akt activity, but area 2 displayed low ERK activity, while area 3 displayed pulsatile ERK activity. HRG (area 4) induced packs of sharp ERK activity pulses enveloped by wide Akt pulses of high amplitude. HGF (area 5) induced highly pulsatile, sharp, synchronous ERK and Akt pulses. However, the large overlap of HGF with area 1 indicates that many cells had adapted and behaved as starved cells. The ErbB1 ligands (BTC - area 6, EGF - area 7, EPR - area 8)^12^ induced similar dynamics consisting of synchronous ERK/Akt pulses. The relative width of the ERK/Akt pulses, however, varied from sharp (BTC) to wide (EPR) as well as plateau-like (EGF). Consistently with a west-east t-SNE projection axis, HRG and IGF displayed higher Akt, while BTC, EGF and EPR displayed lower Akt amplitudes.

A limitation of examining top prototypes is that they might not faithfully reflect the heterogeneity of signaling states in a class. For example, this is illustrated by the absence of top prototypes from areas where GFs overlap (Fig.1b). Therefore, we used an alternative sampling strategy to identify prototype trajectories, whose CNN features are as uncorrelated as possible (Supplementary Fig.S3b,c). To ensure that the selected trajectories are still representative of each class, only trajectories that reached a minimal threshold of prediction confidence for their actual class were considered. This resulted in a better coverage of trajectories in the CNN feature space, while maintaining class specificity. Visual evaluation of prototype trajectories sampled by different methods provides a more complete intuition about discriminant features that characterize the classes. It is also of interest to identify trajectories for which the CNN prediction was wrong despite exhibiting a large confidence (Supplementary Fig.S3d). Indeed, the ErbB1 ligands BTC, EPR and EGF that share trajectories with similar features, were often wrongly classified as another ErbB1 ligand. The comparison of such prototypes might help to comprehend how different receptors, of the same family, are differently wired to downstream signaling networks and give rise to a range of signaling states.

The dynamic information carried by signaling pathways often relies on local signal shapes such as the steepness of pulses, the time interval between pulses^5,6,13^ or the decay kinetics^14^. Using CAMs, we identified minimal signaling motifs that discriminate trajectories induced by each ligand. We then used DTW shape-based clustering to evaluate their distribution within different GF classes (Fig.2a,b). This again provided intuition about GF-specific signaling dynamics: 1. The ErbB1 ligands BTC, EGF, EPR all displayed a mix of synchronous ERK/Akt pulses in which Akt amplitude was lower than ERK amplitude (clusters 2, 3), or wider ERK/Akt pulses with a very low Akt amplitude (cluster 4); 2. HRG led to a peculiar pattern consisting of multiple sharp ERK pulses under larger Akt pulses (cluster 5); 3. IGF displayed sustained Akt, and baseline ERK activity with occasional pulses (clusters 6, 7). The identification of such minimal GF-dependent signaling motifs might provide valuable insights for modelling the networks that produce these dynamic states^15^.

**Figure 2:**
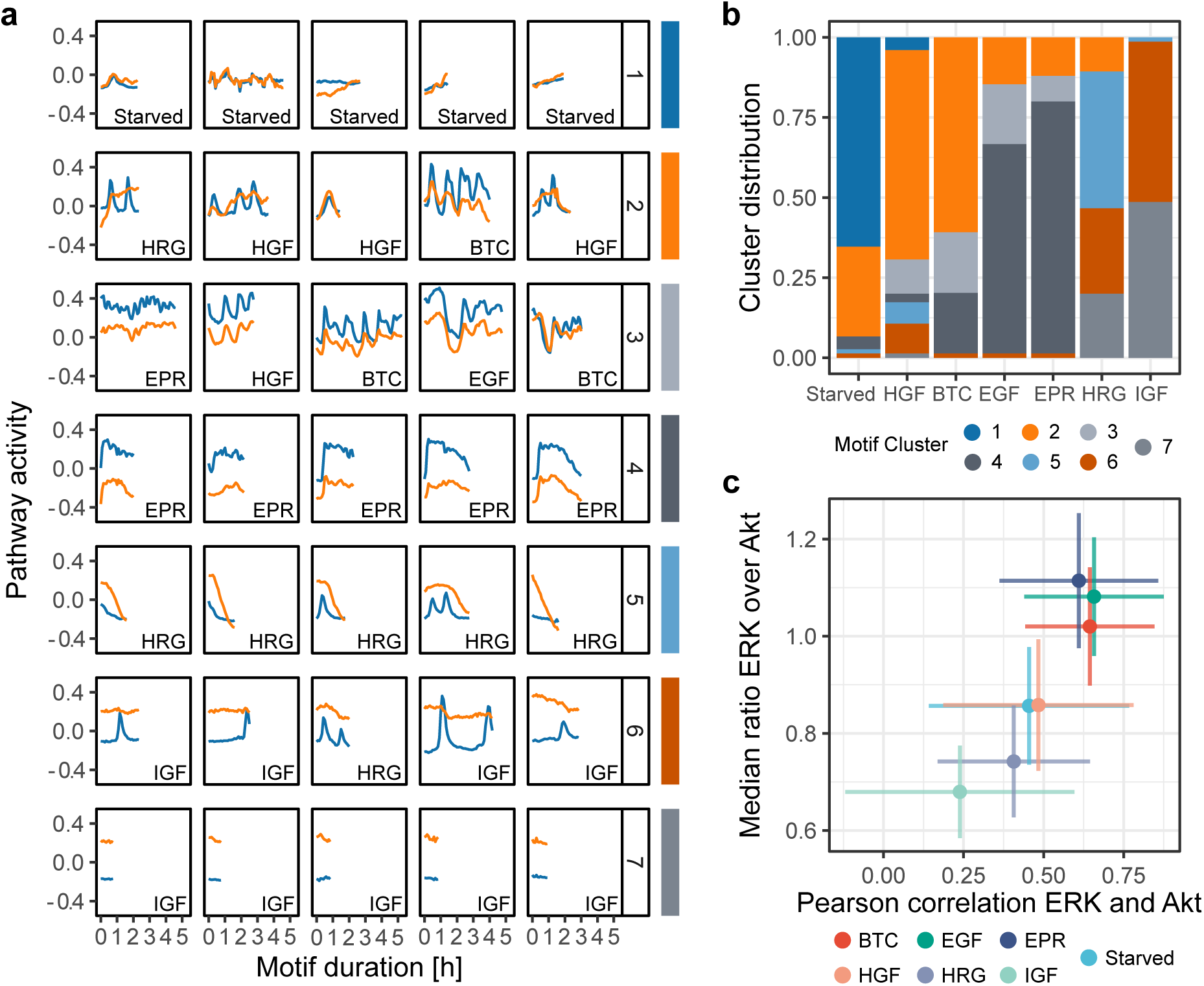
Discriminative motifs in ERK/Akt trajectories highlight GF signaling signatures and ease the identification of interpretable, GF-specific features. **a**, Discriminative signaling motifs were extracted from the training and validation top prototypes using a CAM-based approach (see Methods). The motifs were clustered using DTW distance and Ward’s linkage. Representative motifs of each cluster, based on the minimization of median intra-cluster distance, are displayed. Bottom-right labels indicate the class of the trajectory from which the motif was extracted. Each pathway activity is preprocessed by removing its respective average in the training set. **b**, Distribution of the signaling motifs clusters across the GF treatments. **c**, Scatterplot of Pearson correlation coefficient between ERK and Akt trajectories against median ratio of ERK over Akt activity. For each trajectory, ratios are computed on raw data, at each time point and summarized with median. Crosses indicate the mean values and the standard deviations of all raw single-cell trajectories.

The three components of CODEX, i.e. projection of CNN features, identification of prototypes, and motif extraction, provided intuitive insights that allowed us to empirically identify combinations of interpretable features able to identify different signaling states. For example, CODEX suggested that the synchrony between ERK and Akt as well as their amplitude ratio were highly discriminative across GF stimulations. We verified this by computing the temporal correlation between ERK and Akt as well as the median ratio of ERK over Akt (Fig.2c). The frequency of ERK and Akt pulses and their synchrony were also suggested and validated as discriminative features (Supplementary Fig.S4).

To showcase the versatility of CODEX and benchmark it against classical approaches, we analyzed additional biosensor signaling datasets from previous studies. In one example, we analyzed SMAD2 dynamics in a TGFβ dose response^7^. Each TGFβ dose induced different, heterogeneous SMAD2 dynamic trends that were isolated by shape-based clustering using DTW distance. CODEX identified almost identical prototype states with a clustering of the CNN features (Supplementary Fig.S5 and Note 3). In another example, we analyzed a p53 dynamics dataset consisting of 12 different cancer cell lines subjected to 5 different doses of ionizing radiation. The combination of cell lines and radiations doses yields a total of 60 classes with dozens of p53 trajectories each^10^. The size of this dataset poses a major challenge for data visualization and mining. CODEX allowed us to rapidly identify discrete p53 dynamic states and evaluate their distribution across conditions. This hinted that cell line identity rather than radiation dose dictates dynamic p53 states, and successfully recapitulated previously identified cell line-specific signaling states (Supplementary Fig.S6 and Note 4). Finally, we used CODEX on a time-series dataset representing the movement speed of male and female *Drosophila Melanogaster* under day and night light conditions^16^ (Supplementary Fig.S7 and Note 5). The trajectories in this dataset are significantly different in length and shapes compared to the other datasets. Despite this, with the same CNN architecture, the model converged to an excellent classifier, whose output correlates with interpretable features that were previously reported.

CODEX revisits conventional pipelines for extracting and mining features from large datasets. Instead of centering the analysis around predefined features and their statistical significance, our approach learns features and uses them to highlight important pieces of data. We can therefore dissect complex time-series datasets, identify trends in signaling dynamics and confront them directly against the data. The combination of low-dimensional projection of large time-series datasets, identification of prototypes and local motifs makes it a valuable tool for early data exploration. We have shown that CODEX provides a universal approach to quickly build hypotheses and identify phenotypes in dynamical readouts from a wide variety of biological systems. Beyond this, CODEX demonstrates how modern machine learning models, often criticized for their opacity, can reduce the workload of mining large datasets and propose targeted, interpretable analysis.

## Methods

### Cell culture and biosensor imaging

MCF10A cells were cultured in DMEM:F12, 5% horse serum, 20 ng/ml recombinant hEGF (Peprotech), 10 µg/ml insulin (Sigma-Aldrich/Merck), 0.5 mg/ml hydrocortisone (Sigma-Aldrich/Merck), 200 U/ml penicillin and 200 µg/ml streptomycin. GFs stimulation experiments were executed after 2 days’ starvation in DMEM:F12, 0.3% BSA (Sigma-Aldrich/Merck), 0.5 mg/ml hydrocortisone (Sigma-Aldrich/Merck), 200 U/ml penicillin and 200 µg/ml streptomycin. hEGF, BTC, EPR, HGF, HRG and IGF (Peprotech) were pre diluted in the starving medium and added to cells under the microscope.

H2B-miRFP703, ERK-KTR-mTurquoise2 and FoxO3a-mNeonGreen constructs were generated and subcloned in the piggy PiggyBac vectors pMP-PB, pSB-HPB and pPB3.0.Blast as previously described^17^. Upon transfection of these plasmids with FuGene (Promega), cells were treated with 2.5 µg/ml Puromycin, 25 µg/ml Hygromycin and 5 µg/ml Blasticidin to select stably expressing cells. To achieve uniform biosensor experiments cells were further cloned. For imaging experiments, MCF10A cells were seeded on 5 µg/ml Fibronectin (PanReac AppliChem) on 24 well 1.5 glass bottom plates (Cellvis) at 1×105 cells/well density two days before the experiment. Time-lapse epifluorescence imaging was executed with an Eclipse Ti inverted fluorescence microscope (Nikon) equipped with a Plan Apo air 40× (NA 0.9) objective. Images were acquired with a 16-bit Andor Zyla 4.2 plus camera and with the following excitation and emission filters (Chroma): far red: 640nm, ET705/72m; NeonGreen: 508nm, ET605/52; mTurquoise2: 440nm, HQ480/40. Images were acquired with 1024×1024 resolution with 2×2 binning.

### Automated image analysis

To obtain single-cell bivariate signalling trajectories of ERK and Akt activities we used a dedicated image analysis pipeline, as previously described^17^. First, we trained a random forest classifier based on different pixel features (Ilastik^18^) to separate H2B-miRFP703 fluorescence from background signal. The 32-bit nuclear probability channel produced by pixel classification was then used for nuclear segmentation with CellProfiler 3.0^19^. A 7 pixels expansion of nuclear segmentation with 2 pixels separation was used to produce a ring-shaped ROI in the cytosol. A ratio of the median pixel intensities of the cytosol over the nucleus masks was then calculated. Centroid-based single-cell tracking was executed with MATLAB using µ-track 2.2.1^20^.

ERK and Akt activities were calculated as cytosol/nuclear (C/N) ratio of average pixel intensities in cytosol and nuclear ROIs in the respective fluorescence channels. Color coded images of ERK and Akt activities (fig. S2B and C) were generated by color coding nuclear segmentation with the C/N values for each cell in each time point (CellProfiler 3.0).

ERK and Akt data analysis in single-cell trajectories was carried out with custom R codes. Heatmaps of signalling trajectories (fig. S2D) and average plus 95% confidence interval (fig. S2E) were generated with Time Course Inspector^21^.

### CNN architecture, training parameters, data augmentation and preprocessing

All CNNs were built on the same fully convolutional architecture^11^, only the number of filters in the last convolutional layer, *i.e*. the number of CNN features used for classification and projection, was varying (Supplementary Table S1). Batch normalization and rectified linear units (ReLU) activation were used after each convolution layer. This standard setup was shown to improve speed of training, help training from scratch and prevent overfitting^22,23^. In this configuration, convolutions on bivariate data are done as if the signal were an image with 2 rows of pixels and a single color channel. All models were trained to minimize cross-entropy loss, with L2 regularization of weight 1e-3. Learning rates were initialized at 1e-2 and progressively reduced through epochs. The number of epochs varied from about 20 epochs for the synthetic set to a few hundred for the GF dataset, but all were trained in less than an hour on a consumer-grade GPU (Nvidia RTX 2080 Ti).

For all datasets, input trajectories were preprocessed before being passed to the CNN by subtracting from each channel its average value in the training set. The chosen CNN architecture imposes a fixed input size. We propose to take advantage of this limitation to perform data augmentation by randomly cropping trajectories before presenting them to the network (Supplementary Table S1). The procedure is analogous to what is commonly done on images, where cropping was shown to be efficient at enforcing space-(in our case, time-) invariant features learning^24^. For the GF dataset analysis, we fixed a set of input trajectories to get rid of any variation due to the random crop before creating all figures related to these data.

### Prototype trajectories selection

We use the classification confidence of CNNs to identify prototype curves that are representative of the input classes. The classification output of a CNN consists in a one-dimensional vector of real numbers, in which each number represents a single class. These numbers are not bound to a specific range but a higher number, relative to the rest of the output vector, is a stronger indication that the input belongs to a given class. As is usually performed, we transformed these output vectors with the softmax function. This squeezes all numbers between 0 and 1 and ensures that their sum is equal to 1. Hence, it gives a probabilistic interpretation to the CNN output. We define the classification confidence of a model for a given input as the predicted probability of this input for a given class. We distinguish 2 types of prototypes trajectories. On one hand, the “top prototypes”, which are the trajectories for which the model prediction is correct and for which its confidence is the highest in a set of input trajectories. On the other hand, the “uncorrelated prototypes” which is a subset of input trajectories that we extract in two steps. First, the set of input trajectories is filtered to retain those for which the model confidence in the correct prediction reaches a predefined threshold. Second, a greedy algorithm chooses one by one trajectories in this filtered set such that their CNN features are as least correlated as possible between each other. This procedure is initiated by selecting the trajectory which has the highest median Pearson correlation to all the other trajectories in the filtered set.

For the GF dataset (Fig.1b,c), the 10 top prototypes from each class in the validation set were selected. For the synthetic dataset (Supplementary Fig.S1b), 8 uncorrelated prototypes with minimal confidence of 90% were chosen in the training and validation sets pooled together. For the drosophila movement dataset (Supplementary Fig.S7b), the top prototypes for each class were chosen in the training and validation sets pooled together.

### Motif mining and clustering with CAMs

Class-discriminative motifs were identified with Class-Activation Maps (CAMs), a technique to reveal class-specific regions according to a CNN classifier in the input trajectories^11^. CAMs assign a quantitative value to each data point in input trajectories, large values indicate points that largely affect the model prediction towards a class of interest.

Here, we define a motif as a continuous stretch of points in the input trajectories that are recognized as important by the CNN for a given class, as indicated by CAMs. This approach is analogous to what was already proposed in computer vision^25^. To obtain the motifs, the points in an input trajectory are first classified as “relevant” or “non-relevant” for the class by binarizing CAM values using Li’s minimum cross entropy threshold^26^. This results in a collection of continuous “relevant” segments which are expanded by a defined number of points. This helps to better capture the context around a motif and to correct for single points detected as “non-relevant” by the thresholding. This collection of extended segments in a trajectory forms the collection of class-specific motifs in the trajectory.

In order to go beyond the identification of motifs among single trajectories, we established a motif mining pipeline to investigate and characterize motifs at the dataset scale. To do so, we first isolate motifs in prototype trajectories from every class in the datasets. The CAMs to identify these motifs are generated towards the actual class of each trajectory. Then, from each trajectory only the longest motif is retained. Finally, all the motifs are compared with dynamic time warping (DTW) distance and partitioned with hierarchical clustering.

For the GF (resp. Synthetic) dataset, 75 (resp. 125) top prototype trajectories (resp. uncorrelated prototype trajectories with minimal confidence of 90%) were selected from both the training and the validation sets to extract the patterns. Motifs were extended by 2 (resp. 0) points and only the longest motif was retained in each trajectory. The motifs were finally filtered to retain motifs shorter than 100 points (resp. longer than 5 points) (Fig.2a, Supplementary Fig.S1c).

### Dynamic Time Warping

Dynamic Time Warping (DTW) distances between CAM-motifs were computed with the R package *parallelDist*, with the step pattern “symmetric2”. Pairwise distances were normalized by the sum of the lengths of both motifs of the pair.

### Medoids and centroids for motifs clusters and CNN features clusters

To present the content of the motif clusters (Fig.2a, Supplementary Fig.S1c), sets of representative motifs were selected. The selection process explicitly relies on the distance matrices that were used to perform the clustering. Namely, the medoid motifs are the motifs for which the median DTW distance to all other motifs in the same clusters is minimal.

A similar procedure was followed to choose representative trajectories from CNN features clusters (Supplementary Fig.S5f right column, S6d). These clusters were defined by hierarchical clustering using the L1 distances between the CNN features of the trajectories. This distance matrix was used to select trajectories that minimized intra-cluster distance. For the TGFβ/SMAD2 dataset, centroids were used in place of medoids, *i.e*. trajectories that minimized the average intra-cluster distance (and not the median) were selected.

### t-SNE projections

The t-SNE projections of the CNN features were performed with the implementation in the Python library *sklearn*.

### Peak Detection

The number of ERK/Akt activity peaks were calculated with a custom algorithm that detects local maxima in time series. First, we applied a short median filter to smoothen the trajectories. Then, we ran a long median filter to estimate the long-term bias, which was then subtracted from the smoothed time series. We only kept the positive values. These smoothed and detrended time series were then rescaled to [0,1]. Finally, peaks were detected as points that exceeded a threshold that we manually set at 0.12 for ERK and 0.10 for Akt.

### Synthetic data

Synthetic data were created by generating trajectories that always comprise 4 events of pulses (Supplementary Fig.S1). Each pulse can be of two types: either a full Gaussian peak or a Gaussian peak truncated at its maximum. The side of the peak being truncated is random for each peak. The equation to generate a single peak event is:

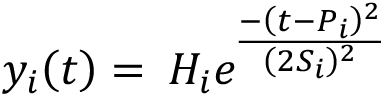

Where 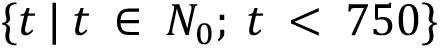 is the discrete time variable, stopping when the desired length of trajectory is reached (here 750); P is the discrete random variable for the time of event occurrence, follows *U*{0, 750}; H is the height of the peak, follows *U*(1, 1.5) and S relates the width of the peak, follows *U*(15,25). After truncation of a peak, the final equation for a single peak is:

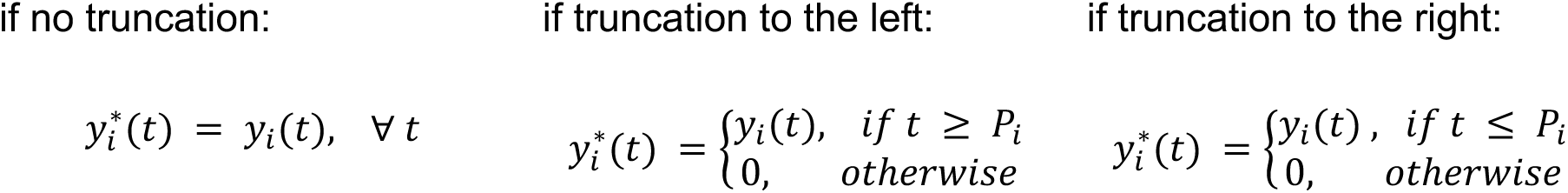

The number of truncation events for a single trajectory is drawn from *U*{0, 1, 2}for trajectories of the first class and *U*{2,3,4}for trajectories of the second class. Finally, the whole trajectory is obtained by summing all independent peak trajectories:

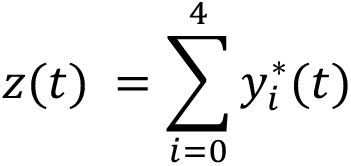

We add the hard constraint that each peak (i.e. each realization of P in a trajectory) must be at least 75 time points away from each other.

The final synthetic dataset contains 10 000 trajectories in each class, from which 70% were used for training and the rest for validation.

## Code availability

The code necessary to run CODEX, along with user-friendly Jupyter notebooks and the interactive application to browse the t-SNE projection of CNN features are freely available at: https://github.com/majpark21/CODEX.

## Data availability

All data and markdowns to reproduce every figure can be downloaded from: https://doi.org/10.17632/4vnndy59fp

## Acknowledgements

The authors thank A. Loewer for providing the TGFβ/SMAD2 dataset as well as J. Stewart-Ornstein and G. Lahav for providing the p53 dataset.

## Authors contribution

M-AJ, MD, RS and OP designed the study. M-AJ wrote the source code and analyzed the data. PAG performed biosensor imaging. M-AJ, OP, PAG and MD wrote the manuscript. OP, MD and RS supervised the project.

## Supplementary Information

**Figure S1:**
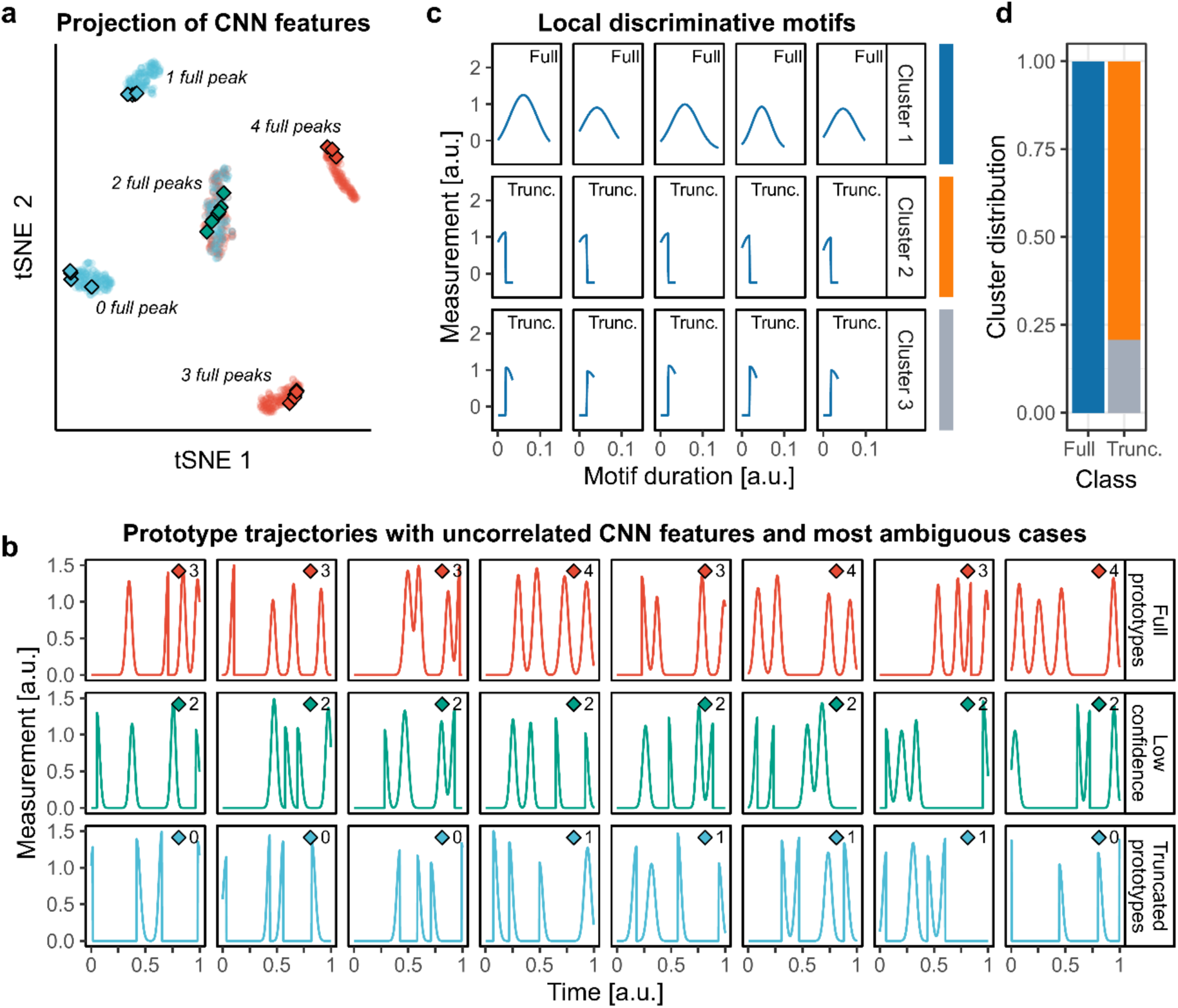
CNN highlights discriminative dynamics motifs in synthetic data. **A** synthetic dataset resembling pulsatile signaling activities was created and a CNN was trained to recognize its 2 classes (see Methods and Supplementary Note 2). Each trajectory comprises 4 pulsing events that consist of either full Gaussian peaks or half-truncated Gaussian peaks. Events are triggered at random locations but with a minimal distance from each other. The class labeled Full displays predominantly full peaks (2, 3 or 4), while the class labeled Truncated displays predominantly truncated Gaussian peaks (0, 1 or 2 full peaks). **a**, t-SNE projection of the CNN features of the trajectories in the validation set. Each cluster of points regroups trajectories based on their number of full peaks. Diamonds indicate prototype trajectories. **b**, Prototype trajectories (blue and red) sampled by selecting trajectories whose CNN features were uncorrelated (see Methods), and trajectories for which the model confidence is lowest (green). Trajectories are the ones indicated with diamonds in (**a**). The number of full peaks in each trajectory is indicated next to the diamond symbol. The minimal threshold of model confidence to sample prototypes was set to 90%. **c**, Motifs extracted with the CAM method (see Methods). Motifs were clustered with DTW distance and Ward linkage. Representative medoid motifs of the clusters are shown. The class of the trajectory from which the motif was extracted is indicated in the top-right corner. **d**, Distribution of the motifs in the clusters according to the class of their origin trajectory.

**Figure S2:**
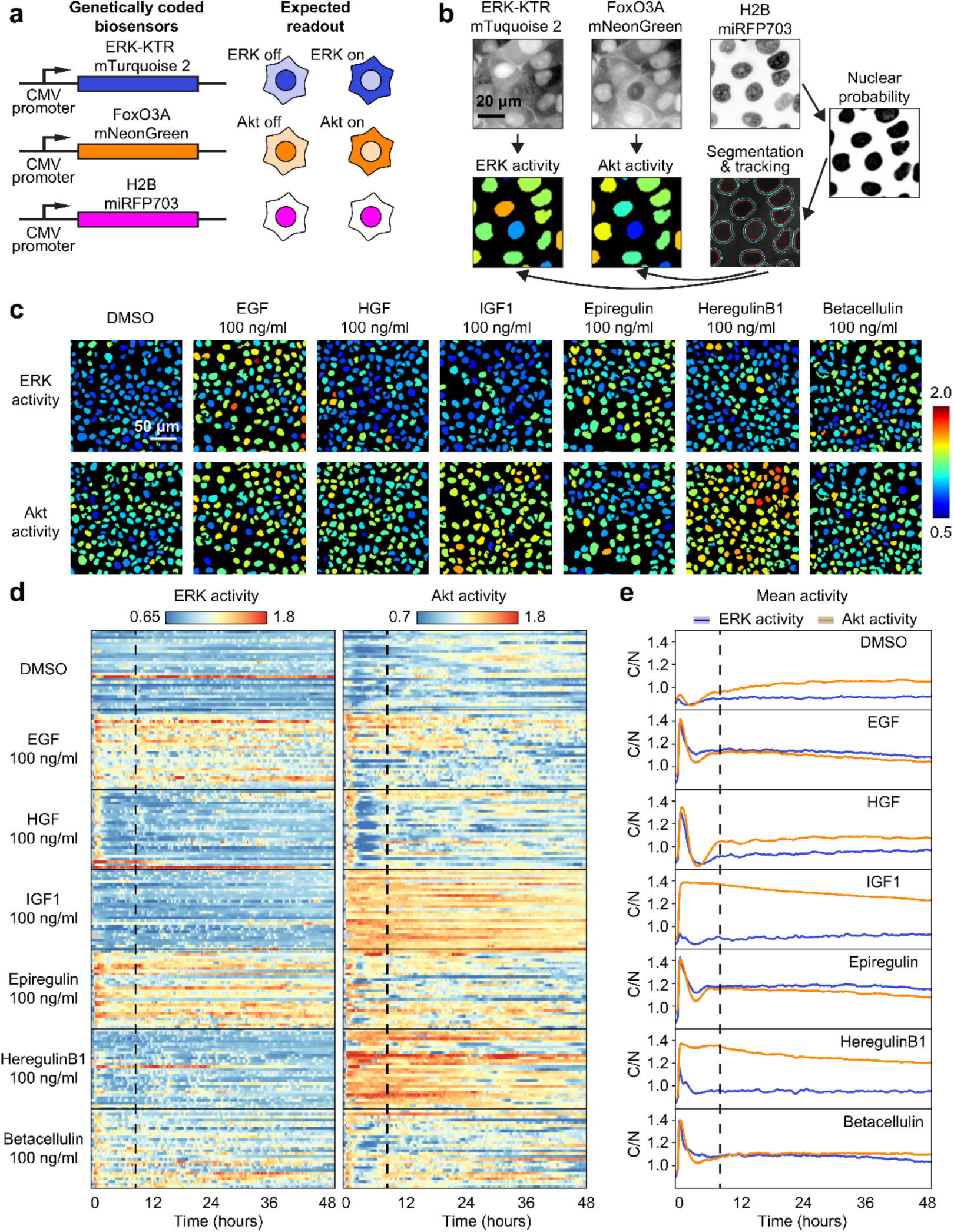
Fluorescent biosensors of ERK and Akt activity report single-cell signaling dynamics in response to growth factors. **a**, Scheme of the multiplexed, genetically-encoded ERK (ERK-KTR-mTurquoise2) and Akt (FoxO3A-mNeonGreen) biosensors and the constitutive nuclear marker (H2B-miRFP703). Both biosensors translocate from nucleus to cytosol when phosphorylated by their respective kinase. **b**, Image analysis pipeline to obtain single-cell ERK/Akt trajectories. A random forest classifier (Ilastik) was trained to segment nuclei. Nuclei (red outline) are used for tracking each cell across the time-lapse. Expansion of the nuclear mask provides a cytosol area (cyan outline). The ratio of pixel intensities of the cytosolic over the nuclear area provides a proxy for ERK or Akt activity, which is displayed over each nuclei with a specific color code (high/low ERK/Akt activity = warm/cold colors respectively). **c**, ERK and Akt activity channels in MCF10A monolayers 25 hours after the stimulation with the different growth factors. **d**, Randomly selected ERK and Akt activity single-cell trajectories from MCF10A monolayers treated with the different growth factors. The dashed black vertical line represents the first frame of the portion of signaling trajectories used to train CNN. GFs have been added at the beginning of the traces. **e**, Population averages of ERK/Akt activities of a whole field of view.

**Figure S3:**
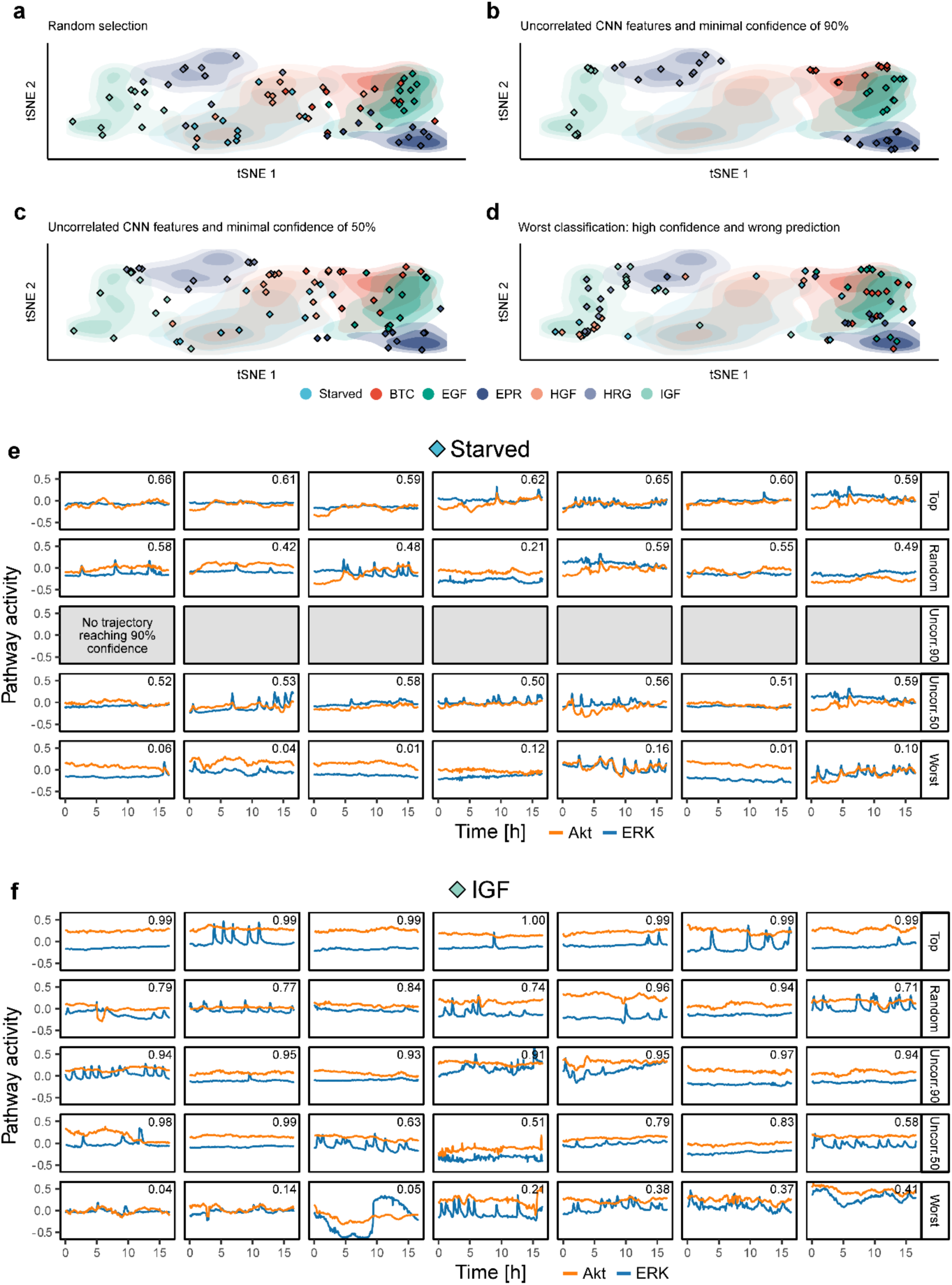
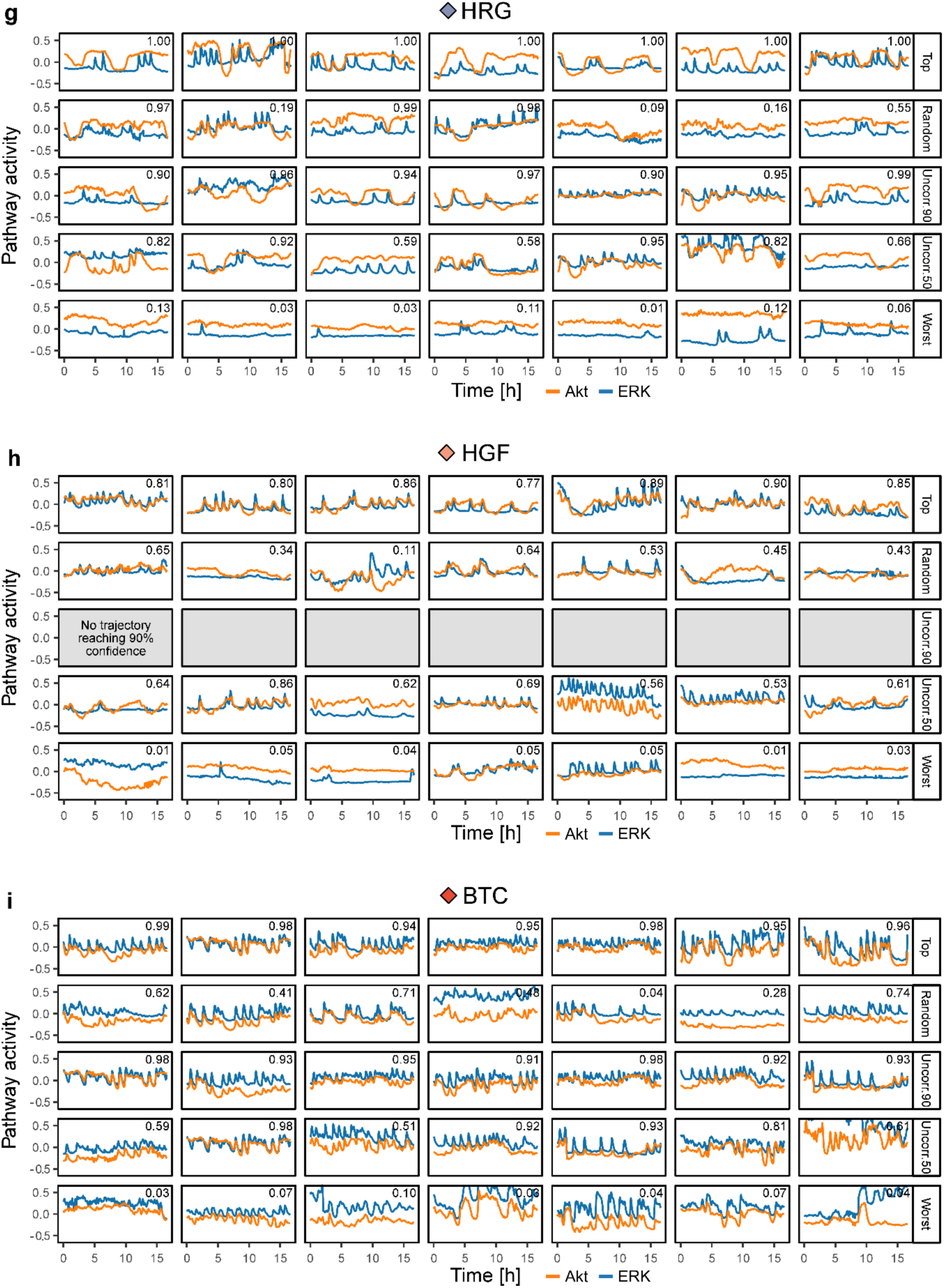
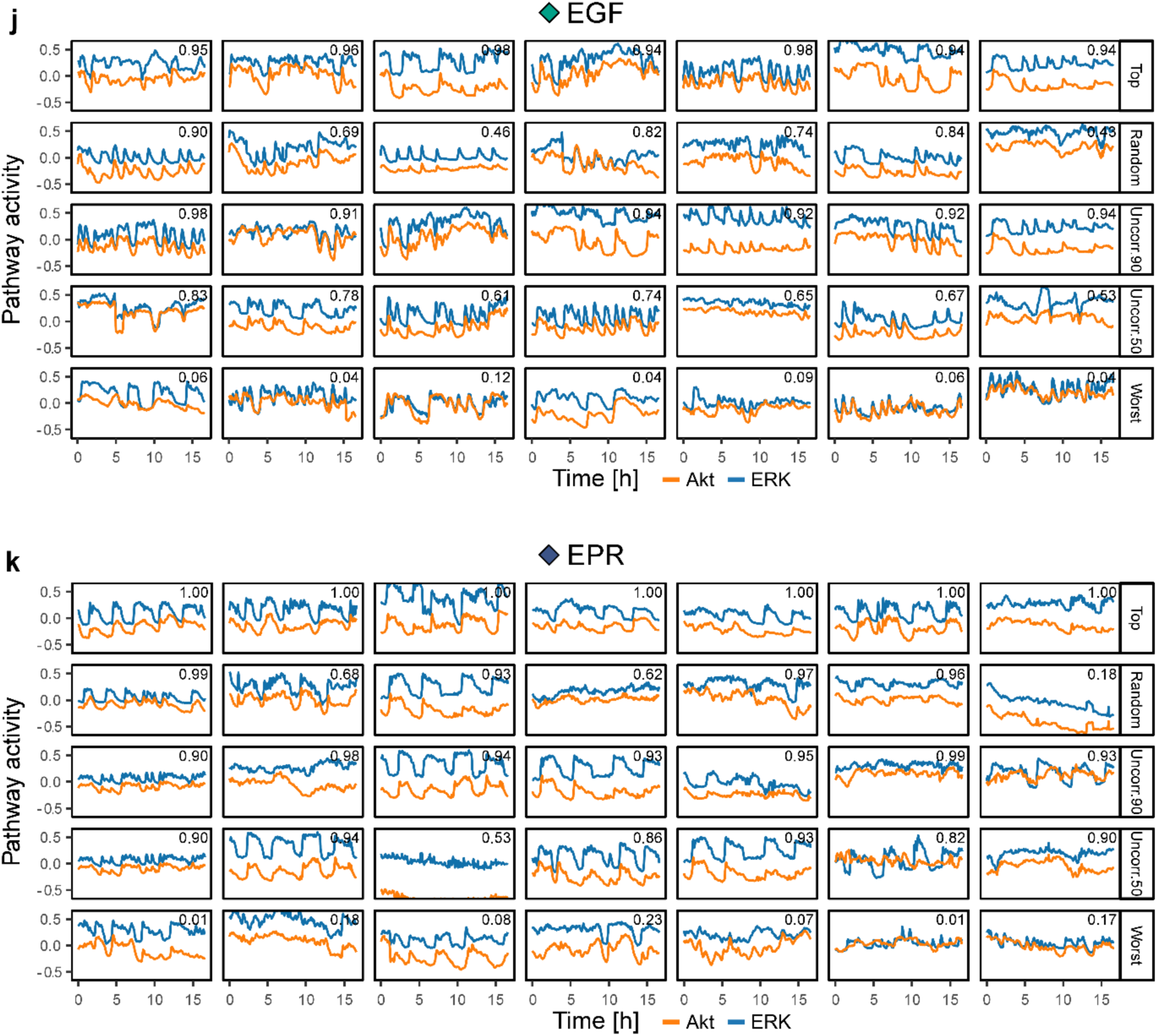
Comparison of prototype trajectories sampling strategies. **a-d**, Different strategies to sample prototype trajectories from the GF ERK/Akt dynamics dataset are presented on the same t-SNE projection of the CNN latent features of trajectories from the validation set. Prototype trajectories are indicated with diamonds, 10 trajectories are shown for each GF with each strategy. Missing diamonds in **b** or **c** indicate that none, or only a small proportion of trajectories reached the minimum confidence threshold. **a**, Random selection, **b**, Uncorrelated CNN features and minimal confidence of 90%, **c**, Uncorrelated CNN features and minimal confidence of 50%, **d**, Worst classification: high confidence and wrong prediction. **e-k**, Comparison of prototype trajectories identified using different sampling strategies. **e**, Starved, **f**, IGF, **g**, HRG, **h**, HGF, **i**, BTC, **j**, EGF, **k**, EPR.

**Figure S4:**
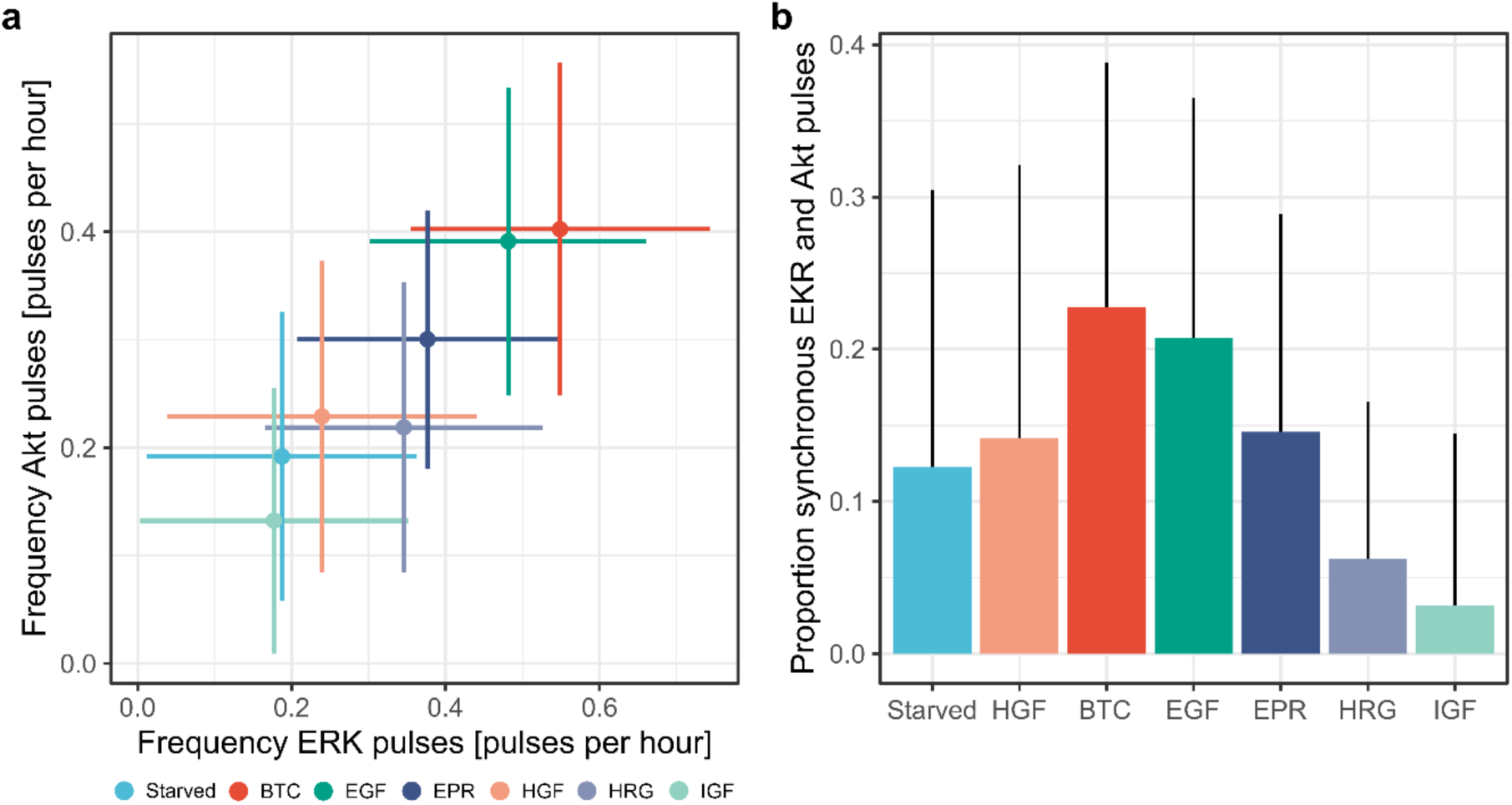
Quantification of additional interpretable, class-discriminative features identified by visual inspection of the CODEX output. **a**, Scatterplot of frequency of ERK peaks versus frequency of Akt peaks in single-cells. Crosses indicate the mean values and the standard deviations. Colors indicate the GF and are the same as in (**b**). **b**, Proportion of synchronous ERK and Akt peaks in single-cells. Mean and standard deviation are shown.

**Figure S5:**
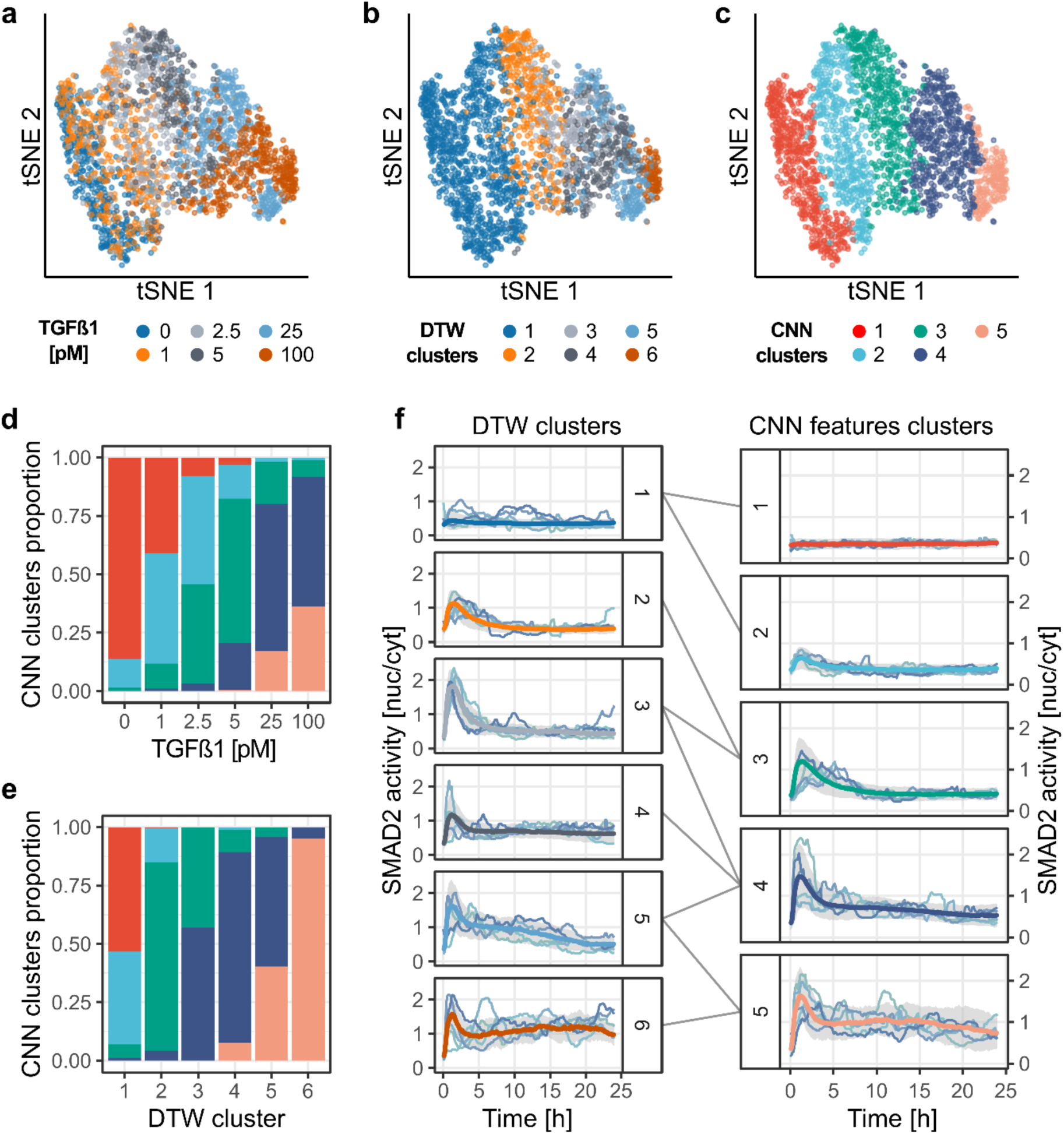
CODEX identifies TGFβ dose-dependent signaling states. MCF10A cells were exposed to a TGFβ dose response and single-cell SMAD2 responses were recorded using a fluorescent biosensor. A CNN was trained to classify SMAD2 activity trajectories according to the TGFβ dose. Trajectories were directly clustered based on their CNN features vector and the results were compared to a constrained DTW clustering (see Methods and Supplementary Note 3). The SMAD2 trajectories and the DTW clusters were provided by the original authors^7^. L1 distance was used on raw CNN features and trajectories partitioned with hierarchical clustering and Ward linkage. **a-c**, t-SNE projection of the CNN latent feature space for the training, validation and test sets pooled together. Trajectories representations are colored according to: the TGFβ dose (**a**), DTW clusters (**b**) or CNN features clusters (**c**). **d-e**, Distribution of the trajectories in the CNN features clusters according to their corresponding TGFβ dose (**d**) and their DTW cluster (**e**). Same colors as in (**c**). **f**, Comparison of DTW clusters and CNN features clusters. Median trajectories for each cluster are reported in bold colored lines, colors matching (**b**) and (**c**), grey shade indicates interquartile range. A random sample of trajectories for DTW clusters, as well as the centroid trajectories for each CNN-features cluster are shown.

**Figure S6:**
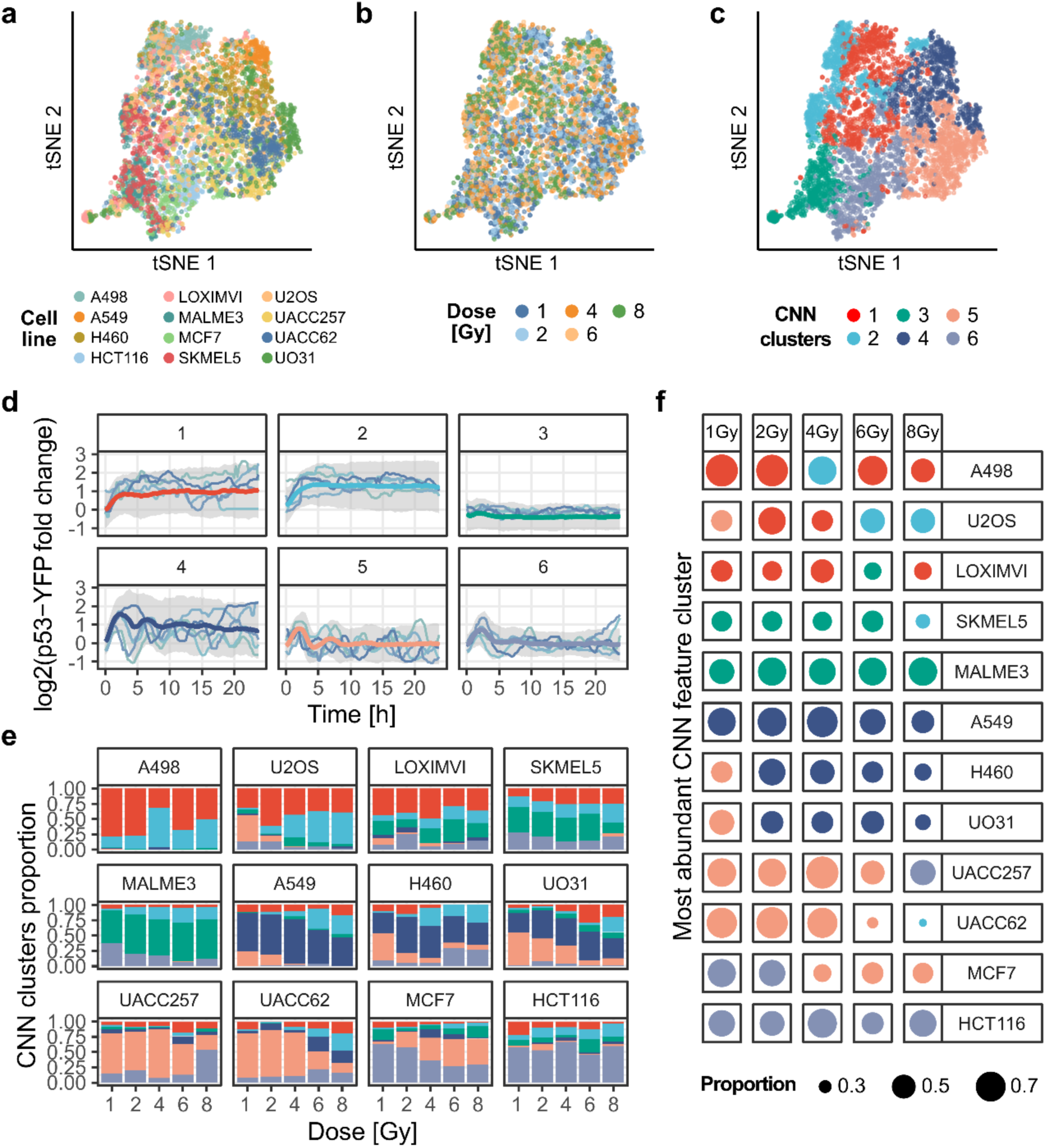
CODEX identifies cell line-specific p53 responses under increasing ionizing radiation doses. 12 different cell lines were exposed to 5 doses of ionizing radiations and single-cell p53 responses were reported using a fluorescent reporter. A CNN was trained to recognize the combination of cell line and radiation dose from p53 trajectories (i.e. one out of 60 classes; see Methods and Supplementary Note 4). The p53 trajectories were provided by the original authors^10^ and were clustered according to their standardized CNN features with L1 distance and partitioned with hierarchical clustering and Ward linkage. **a-c**, t-SNE projection of the CNN features for the training, validation and sets pooled together. Trajectories representation are colored according to: the cell lines (**a**), ionizing radiation doses (**b**), or CNN features clusters (**c**). **d**, Representative p53 trajectories of the clusters identified using CNN features clustering. Median trajectories are reported in bold colored lines, colors matching (**c**), grey shade indicates interquartile range. Individual traces indicate the medoid p53 single-cell trajectories for each cluster. **e**, Distribution of the trajectories in the CNN feature clusters according to cell lines and ionizing radiation doses. Same colors as in (**c**). **f**, Most abundant CNN features cluster for all trajectories in each combination of cell line and radiation dose combinations. Color of the dot indicates the most abundant cluster; size of the dot indicates the proportion of cells classified in the indicated cluster. Same colors as in (**c**)

**Figure S7:**
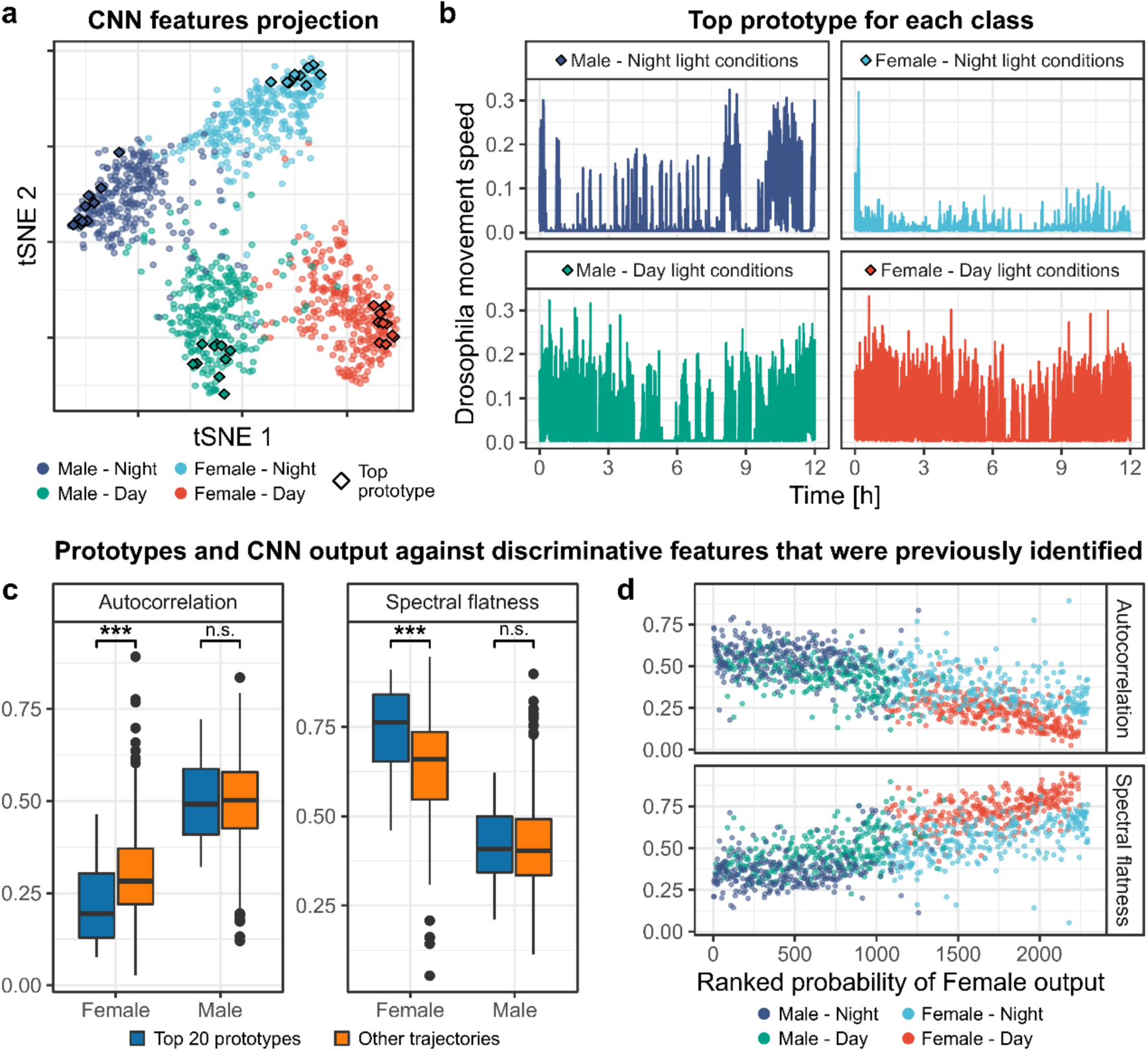
CODEX identifies different sex- and light-dependent *Drosophila* movements. Male and female *Drosophila* movements in a tube were recorded over 12h, in presence or absence of light, and reported as univariate time-series^16^. A CNN was trained to recognize the combination of sex and light conditions based on the individual movements (see Methods and Supplementary Note 5). **a**, t-SNE projection of the CNN features for the training, validation and test sets pooled together. Diamonds indicate the 10 top prototypes for each class. **b**, The top prototype trajectory for each class. **c**, Comparison of autocorrelation levels and spectral flatness between the 20 top female prototypes and 20 top male prototypes against non-prototype trajectories. Female (resp. male) prototypes were obtained by pooling the 10 top prototypes of female (resp. male) drosophila under both day and night light conditions. Medians of the distributions were compared with Wilcoxon’s test; ***: p-value < 0.01. **d**, Autocorrelation level and spectral flatness against the probability of trajectories to be classified as a female class. The latter probability is the sum of the probability of trajectories to belong to the female-day and female-night classes. The resulting sum was ranked, such that rank 1 corresponds to the probability of the trajectory that has the lowest prediction outcome to be classified as female. Autocorrelation levels were computed by averaging autocorrelation values for lags ranging from 1 to 10 time points.

## Supplementary Video

**Supplementary Video S1: Interactive application to project and browse CNN features with t-SNE.** The video presents the interface of CODEX’s companion application to visualize single-cell trajectories in a t-SNE projection of their CNN features. A projection of the CNN features learnt for the GF validation dataset was chosen as illustrative example. The CNN that builds the features and the t-SNE parameters are identical to the ones used to generate Fig.1b. The top banner regroups the parameters of the t-SNE projection. The bottom-left panel represents the projection itself, where each point corresponds to a single-cell, bivariate trajectory of ERK and Akt activity. Each point is colored by the GF applied to the corresponding cell. Upon hovering the cursor over a point, the associated trajectory is displayed in the bottom-right panel. In the bottom-right corner, a table displays the CNN prediction probabilities. Trajectories can be overlaid with CAMs and guided backpropagation for any class of the dataset. Both can further be binarized using mean threshold or Li’s minimum cross entropy threshold^26^.

---

**Table S1:** Detailed CNN architectures. All architectures are fully convolutional, each convolution layer is followed by batch normalization and ReLU. Each convolution is performed with suitable padding to retain the size of the input through its successive representations in the CNN. The convolution layers are followed by a global average pooling and a fully connected layer. N feats: number of CNN features, corresponds to the size of the latent representation after global average pooling.

| Input | Classes | N class | Input size | N feats | Convolution layers | Stride | Padding | BatchNorm | ReLU |
| --- | --- | --- | --- | --- | --- | --- | --- | --- | --- |
| Synthetic activity | Full peak abundance | 2 | (1, 750) | 10 | $4 * (5)^{20} - (3)^{20} - (3)^{10}$ | 1 | Same | Yes | Yes |
| ERK, Akt activity | Growth factor treatment | 7 | (2, 200) | 10 | $4 * (3, 5)^{20} - (3, 3)^{20} - (3, 3)^{10}$ | (1, 1) | Same | Yes | Yes |
| SMAD2 activity | TGFβ1 dose | 6 | (1, 270) | 15 | $4 * (5)^{20} - (3)^{20} - (3)^{15}$ | 1 | Same | Yes | Yes |
| p53 activity | Ionizing radiation dose + Cell line | 60 | (1, 75) | 15 | $4 * (5)^{20} - (3)^{20} - (3)^{15}$ | 1 | Same | Yes | Yes |
| Drosophila movement | Male/female + Day/night light | 4 | (1, 4000) | 10 | $4 * (5)^{20} - (3)^{20} - (3)^{10}$ | 1 | Same | Yes | Yes |

( ) kernel size ^ width of layers \* nber identical layers

**Table S2:** Confusion matrix for GF treatments based on ERK/Akt trajectories. Results are from a single run of the test dataset through the classifier.

|  |  | Predicted class |  |  |  |  |  |  |
| --- | --- | --- | --- | --- | --- | --- | --- | --- |
|  |  | BTC | STARVED | EGF | EPR | HGF | HRG | IGF |
| Actual class | BTC | 77 | 10 | 41 | 8 | 10 | 1 | 0 |
|  | STARVED | 4 | 88 | 2 | 1 | 77 | 8 | 6 |
|  | EGF | 29 | 0 | 78 | 8 | 2 | 0 | 1 |
|  | EPR | 8 | 1 | 13 | 97 | 2 | 3 | 0 |
|  | HGF | 5 | 56 | 2 | 0 | 77 | 11 | 2 |
|  | HRG | 1 | 5 | 2 | 0 | 8 | 150 | 11 |
|  | IGF | 0 | 20 | 0 | 1 | 14 | 19 | 171 |

**Table S3:** Classification metrics for GF treatments based on ERK/Akt trajectories. Because of the randomness induced by the random cropping of the input trajectories into segments suitable for the CNN input dimensions, the test set was run 1000 times through the CNN to estimate the metrics. Values indicate the averages and the standard deviations of the metrics.

|  | Precision | Recall | F1 score | Accuracy |
| --- | --- | --- | --- | --- |
| BTC | 53.31 ± 2.11 | 58.50 ± 2.74 | 55.79 ± 2.09 | 65.15 ± 0.81 |
| STARVED | 48.76 ± 1.73 | 50.18 ± 2.28 | 49.45 ± 1.80 | |
| EGF | 64.57 ± 2.11 | 58.00 ± 2.66 | 61.08 ± 2.06 | |
| EPR | 76.41 ± 2.22 | 83.27 ± 2.14 | 79.67 ± 1.68 | |
| HGF | 51.62 ± 2.32 | 40.39 ± 2.30 | 45.30 ± 2.13 | |
| HRG | 82.82 ± 1.64 | 77.46 ± 1.91 | 80.03 ± 1.40 | |
| IGF | 74.58 ± 1.49 | 90.01 ± 1.30 | 81.56 ± 1.13 | |

## Supplementary notes

**Supplementary note 1: Rationale for the choice of CNN architecture.**

We encourage the use of a simple CNN architecture (Supplementary Table S1) that was previously reported in the literature, and that we also found to be a solid baseline through a wide range of datasets^11,27^. We believe that as little effort as possible should go into the network design and training, such that emphasis can be put on the interpretation of results. We thus favorized the ease of model training over predictive power, with smaller models that are easier and faster to train.

This CNN architecture also displays desirable aspects for our application. First, its plain feed-forward structure reduces the range of parameters to tweak if the classification performance is not satisfactory with the defaults. Optimizing a CNN architecture is a very time-consuming activity that requires expertise and, despite recent advances^28,29^ still involves a lot of trials and errors. Reducing the time spent on this step is essential to quickly start with mining the results. Second, the reduced number of parameters enables fast training, even on consumer-grade GPU, and the use of smaller training datasets. The latter implies that training a CNN from scratch is realistic even with relatively small datasets that are usually obtained in single-cell biology. This is an important requirement in cases where no similar and large annotated datasets can be used for transfer learning. Third, a strong countermeasure to overfitting is embedded in this architecture, in the form of a global average pooling (GAP) layer. Not only does GAP help to obtain a reasonable model, the introduction of a bottleneck in the size of the latent features is useful to project them in low dimensions without large distortions. Finally, this architecture is compatible with the creation of CAMs which we use to identify characteristic motifs in the trajectories.

Though the exact architecture of the CNN is flexible, the CAM motif mining approach is restraining the choice of architectures because it explicitly requires the use of global average pooling^11^. This limitation can however be circumvented by replacing CAM with another technique that creates saliency maps such as: grad-CAM, a generalization of CAM which is compatible with more architectures^25^ or guided backpropagation, another popular choice^30^.

**Supplementary Note 2: CODEX creates features that isolates dynamics in synthetic data.**

We first evaluated the ability of a supervised CNN classifier to classify the dynamic landscape in a synthetic time-series dataset, even if this separation is not necessary to perform the classification task perfectly. To do so, we created a synthetic dataset resembling pulsatile signaling activities and trained a CNN to recognize the different synthetic classes (Supplementary Fig.1). All trajectories comprise 4 peak events which can be either full Gaussian peaks or half truncated ones. The dataset comprises 2 classes which differ by their number of full peaks. Trajectories from the first class comprise 0, 1 or 2 full peaks, while trajectories in the second class comprise 2, 3 or 4 full peaks (see Methods). Because the abundance of full peaks is the only difference in the process that generates these curves, there is an ambiguous case when an input trajectory comprises 2 full peaks. Hence the theoretical maximum accuracy for this classification task is 80%, a performance that the CNN reached after a few epochs of training.

The projection of the CNN features learnt for this task shows well-separated trajectory clusters which are grouping together trajectories with a common number of full peaks (Supplementary Fig.S1a, b). The trajectories for which the model was most confused between the 2 classes (“Low confidence”) all harbored 2 full peaks, hence were sampled from the only ambiguous case. This empirical observation is interesting because the model is only optimized to minimize the classification loss, such that the clusters containing 0 or 1 full peak and the clusters containing 3 and 4 full peaks could be merged together without affecting the classification performance. This illustration exposes the core intuition behind CODEX, that CNN features can naturally evolve to capture shapes in the data even without hard constraints. Further, we verified that the motifs captured by CAMs (see Methods) were cleanly isolating class-discriminative motifs (Supplementary Fig.S1c, d). As expected, these motifs contained the tips of the peaks which are either full or truncated, but no flat part of the trajectories which are common to both classes. This also shows that both symmetries of truncation are captured. Finally, one can notice a bias towards the right over the left truncation since cluster 2 is less represented than cluster 3 despite being equally abundant in the trajectories. This last point indicates that the abundance of CAM clusters should be taken with caution. The possible sources of variation are the model training itself as well as the selection of trajectories from which to extract the motifs.

**Supplementary Note 3: CODEX identifies TGFβ dose-dependent signaling states.**

To further showcase CODEX, we used it to analyze a dataset in which SMAD2 activity is measured in response to a dose challenge of TGFβ in MCF10A cells^7^. In this study, a fusion reporter between SMAD2 and YFP provides a readout for SMAD2 activity by computing the relative abundance of the reporter between the nucleus and the cytoplasm. We trained a CNN to recognize the doses of TGFβ that were given to the cells based on their SMAD2 trajectories. In the original study, shape-based clustering of single-cell trajectories using DTW revealed a continuum of SMAD2 heterogeneous signaling states in response to the TGFβ dose response.

We used this already published DTW-based clustering as a global indication of the shape of each trajectory. We then compared how the CNN features for each trajectory were arranged with respect to both the TGFβ dose and the DTW clusters. The projection of the CNN features revealed some entanglements between the TGFβ doses which matched the heterogeneity of SMAD2 signaling at all stimulation doses (Supplementary Fig.S5a). Interestingly, this entanglement seemed largely smoothed out when comparing the CNN features to the DTW clusters (Supplementary Fig.S5b). This again hints that despite the CNN features being learnt with the objective to separate the input classes, they still evolve to capture dynamic trends of the data.

From this observation we hypothesized that directly clustering the CNN features could also provide classes of representative dynamics in the data, similarly to what is done by DTW clustering (Supplementary Fig.S5c). We found the resulting clusters to be in slightly better agreement with the DTW clusters than with the TGFβ doses (Supplementary Fig.S5d, e). To summarize, we found that the CNN features clusters also efficiently captured trajectory profiles (Supplementary Fig.S5f). Visual inspection of representative trajectories suggests that the CNN features performed slightly better in separating flat from weak responders in comparison with the DTW clusters (CNN features clusters 1 and 2; DTW cluster 1).

**Supplementary Note 4: CODEX identifies cell line-specific p53 responses under increasing ionizing radiation doses.**

To further showcase CODEX, we used it to analyze a much larger dataset with more classes. We used a study in which p53 activity dynamics was reported in 12 cancer cell lines under exposure to 5 different doses of ionizing radiation, yielding a total of 60 classes^10^. p53 abundance was reported with a live-cell reporter consisting of a p53-YFP fusion construct.

After training, the projection of the CNN features shows that cell lines, irrespectively of the radiation dose, tend to have trajectories with similar features (Supplementary Fig.S6a, b). This strongly suggests that the cell line, rather than the radiation dose, is the major factor of p53 response variability in this dataset.

We then evaluated whether CODEX could recapitulate cell line-specific behaviors that were previously reported. To do so, we clustered the trajectories on the base of their CNN features into 6 clusters (Supplementary Fig.S6c). Similarly to what we observed for the TGFβ/SMAD2 dataset (Supplementary Fig.S5f), we found that the CNN features clusters capture identifiable trajectory shapes (Supplementary Fig.S6d): sustained activity for cluster 1 and 2, flat baseline for cluster 3, oscillations of large amplitude for cluster 4, oscillations of small amplitude for cluster 5 and a single pulse of activity for cluster 6. We then reported the distributions of these different clusters through the combinations of cell lines and radiation doses (Supplementary Fig.S6e, f). We found similar results to the original study, in which curated features were used to discriminate among dynamic signaling states. For example, MCF7 trajectories strikingly switch from a single pulse (cluster 6) to an oscillatory regime (cluster 5) with increasing radiation doses. Similarly, U2OS cells switched from an oscillatory regime (cluster 5) to a more sustained one (clusters 1 and 2) at high radiation doses. Other cell lines, on the contrary, adopted a consistent behavior through all radiation doses such as HCT116 cells which only display a single pulse of p53 activity (cluster 6) or MALME3 cells which display only low amplitude responses (clusters 3 and 6). CODEX could therefore recapitulate important findings in a large time-series dataset with very little human input and in about one hour for training the model.

**Supplementary Note 5: CODEX identifies discriminative features in *Drosophila Melanogaster* speed movement data.**

We then tested if CODEX can be generalized to completely different time-series datasets. For this purpose, we applied CODEX to an univariate time-series dataset that describes the movement speed of *Drosophila* motility in a tube over 12 hours^16^. There are 4 classes in this dataset that correspond to the combination of the *Drosophila* sex and whether the observation is carried under day or night light conditions. The trajectories are very long and show a strong alternance between resting phases and extreme events that are very different from the smooth series in the signaling datasets. Despite this, using the same CNN architecture, the trained model showed excellent prediction capabilities (accuracy of 90%) that are similar to the ones obtained in the original study with a classifier that uses hundreds of classical time-series features (accuracy of 95%). T-SNE projection of the CNN features clearly separated the 4 classes based on *Drosophila* motility trajectories, and identify prototype trajectories (Supplementary Fig S7a,b).

We then investigated whether a correspondence between CODEX’s output and the classical, interpretable features could be found. We observed that the criteria that separates male from female behaviors were retrieved in the CNN predictions. In the previous study, two of the main reported differences between males and females’ movements are that females present a lower autocorrelation structure and a higher spectral flatness in their movement. We could retrieve this trend in the data and observed that the top prototype trajectories for female classes had a significantly lower autocorrelation and higher spectral flatness than their non-prototype counterparts (Supplementary Fig.S7c). More generally, we observed that a lower autocorrelation structure and a higher spectral flatness correlated with increased prediction odds for female classes (Supplementary Fig.S7d). However, the significant sampling of extreme individuals, regarding both features, was not observed for male prototypes. One could have expected male trajectory prototypes to also be significantly different from the other Features but with the reverse effect than females. This could be due to the fact that the CNN was trained to separate all combinations of sex and night and not only the *Drosophila* sex. This absence of trend calls for caution when making a parallel between the CNN predictions and interpretable features from another set, especially if the former was trained on a different grouping than the one of interest.

